# Insight into the mechanism of H^+^-coupled nucleobase transport

**DOI:** 10.1101/2021.12.20.473561

**Authors:** Jun Weng, Xiaoming Zhou, Pattama Wiriyasermkul, Zhenning Ren, Xiuwen Yan, Kehan Chen, Eva Gil Iturbe, Ming Zhou, Matthias Quick

## Abstract

Members of the nucleobase/ascorbic acid transporter (NAT) gene family are found in all kingdoms of life. In mammals, the concentrative uptake of ascorbic acid (vitamin C) by members of the NAT family is driven by the Na^+^ gradient, while the uptake of nucleobases in bacteria is powered by the H^+^ gradient. Here we report the structure and function PurT_Cp_, a NAT family member from *Colwellia psychrerythraea*. The structure of PurT_Cp_ was determined to 2.80 Å resolution by X-ray crystallography. PurT_Cp_ forms a homodimer and each protomer has 14 transmembrane segments folded into a substrate-binding domain (core domain) and an interface domain (gate domain) A purine base is present in the structure and defines the location of the substrate binding site. Functional studies reveal that PurT_Cp_ transports purines but not pyrimidines, and that purine binding and transport is dependent on the pH. Mutation of a conserved aspartate residue close to the substrate binding site reveals the critical role of this residue in H^+^-dependent transport of purines. Comparison of the PurT_Cp_ structure with transporters of the same structural fold suggests that rigid-body motions of the substrate-binding domain are central for substrate translocation across the membrane.

## Introduction

The nucleobase/ascorbate transporter (NAT) family encompasses proteins that are responsible for the uptake of nucleobases in all kingdoms of life. The NAT family of transporters also mediates uptake of vitamin C (L-ascorbic acid) in vertebrates. Nucleobases and their analogs (e.g., allopurinol, 5-fluorouracil, 6-mercaptopurine, acyclovir) have gained special interest in therapeutic applications as they are used in the treatment of solid tumors, lymphoproliferative diseases, viral infections such as hepatitis and AIDS, and inflammatory diseases such as Crohn’s disease and gout, and as antiparasitic drugs such as trypanocides (1–6). In vertebrates, ascorbic acid, the other substrate of NAT proteins, is central in several vital enzymatic reactions and protects tissues from oxidative damage by scavenging free radicals (7–9).

Intestinal and renal (re)absorption of vitamin C is mediated by the epithelial Na^+^-dependent L-ascorbic acid cotransporter SVeTl (SLe23Al), whereas the homologous SVCT2 (SLC23A2) mediates vitamin e transport in metabolically active cells (e.g., in the adrenal glands, pituitary gland, thymus, corpus luteum, retina and cornea) (10, 11). Furthermore, rat SNBT1 was identified as the first Na^+^-dependent nucleobase transporter in mammals (12).

In contrast to these three mammalian NAT members that mediate Na^+^-coupled transport of L-ascorbic acid and nucleobases, the transport of nucleobases by evolutionary distant eubacteria, archaea, filamentous fungi, plants, insects, and nematodes has been described as being driven by a proton gradient (i.e., H^+^-dependent symport) (1, 13, 14). The crystal structures of the NAT members UraA, the uracil/5-fluorouracil transporter of *E. coli*(15, 16), and UapA, the uric acid/xanthine H^+^ symporter of *Aspergillus nidulans* (17) were obtained in substrate (uracil and xanthine, respectively) bound, inward-open state. These structures have been used as a template for computational studies that propose mechanistic models for the entire NAT family (18, 19). However, limited mechanistic information is available for UraA (15, 20) or the crystalized thermo-stabilized, transport-inactive variant of UapA (17), thereby hampering the interpretation of functional data of NAT proteins in structural context. Here, we present the structure of a bacterial NAT member, PurT of *Colwellia psychrerythraea* (PurT_Cp_) at 2.80 Å resolution that, in conjunction with flux and binding studies, provides insight into the mechanism of transport by this family of transport proteins.

## Results

### Purification and functional characterization of PurT_Cp_

A homolog of human SLC23A1 from the bacterium *Colwellia psychrerythraea* 34H was cloned, expressed and purified (Methods). The protein elutes as a single peak on a size-exclusion chromatography and the elution volume is consistent with PurT_cp_ being a homodimer (Supplementary Fig. 1a). This conclusion is also consistent with a crosslinking study which shows dimer formation (Supplementary Fig. 1b). Dimeric assembly was also reported for UraA (16) and UapA (17).

To assess the activity of the purified, detergent-solubilized SLC23A1 homolog we measured direct binding of radiolabeled purines and pyrimidines with the scintillation proximity assay (SPA) (21). Fig. 1a shows that the purified protein binds the purines guanine, adenine, xanthine, and hypoxanthine, whereas the pyrimidines uracil, thymine, or cytosine did not interact with the candidate protein. To further investigate the substrate specificity of the protein, we performed competition binding assays in which the binding of ^3^H-xanthine and ^3^H-guanine was measured in the presence of a 500-fold concentration excess of non-radiolabeled compounds (Fig. 1b). Consistent with the results shown in Fig. 1a, the non-labeled purines guanine, xanthine, adenine, and hypoxanthine inhibited the binding of ^3^H-xanthine or ^3^H-guanine by ≥ 80 %, suggesting competitive binding to a shared binding site. Similar inhibitory effects were observed with 2-amino-6-bromopurine, 6-bromopurine, 6-thioguanine, allopurinol, isoguanine, and purine. Other purines (caffeine, theobromine, theophylline) reduced binding of ^3^H-guanine or ^3^H-xanthine by ≤ 40 %, whereas the purine nucleotides guanosine or xanthosine, or ascorbic acid, the substrate for the mammalian NAT members, failed to elicit an inhibitory effect. Further elucidation of the binding kinetics for ^3^H-xanthine and ^3^H-guanine with the SPA revealed dissociation constants (*K_d_*) of about 2 μM and 3 μM, respectively, and a nucleobase (xanthine or guanine)-to-protein molar binding ratio of ~1 (Fig. 1c, see also Supplementary Figure 2), revealing the presence of a single nucleobase binding site in the purine transporter of *Colwellia psychrerythraea* 34H that we refer to as PurT_Cp_.

**Figure 1:**
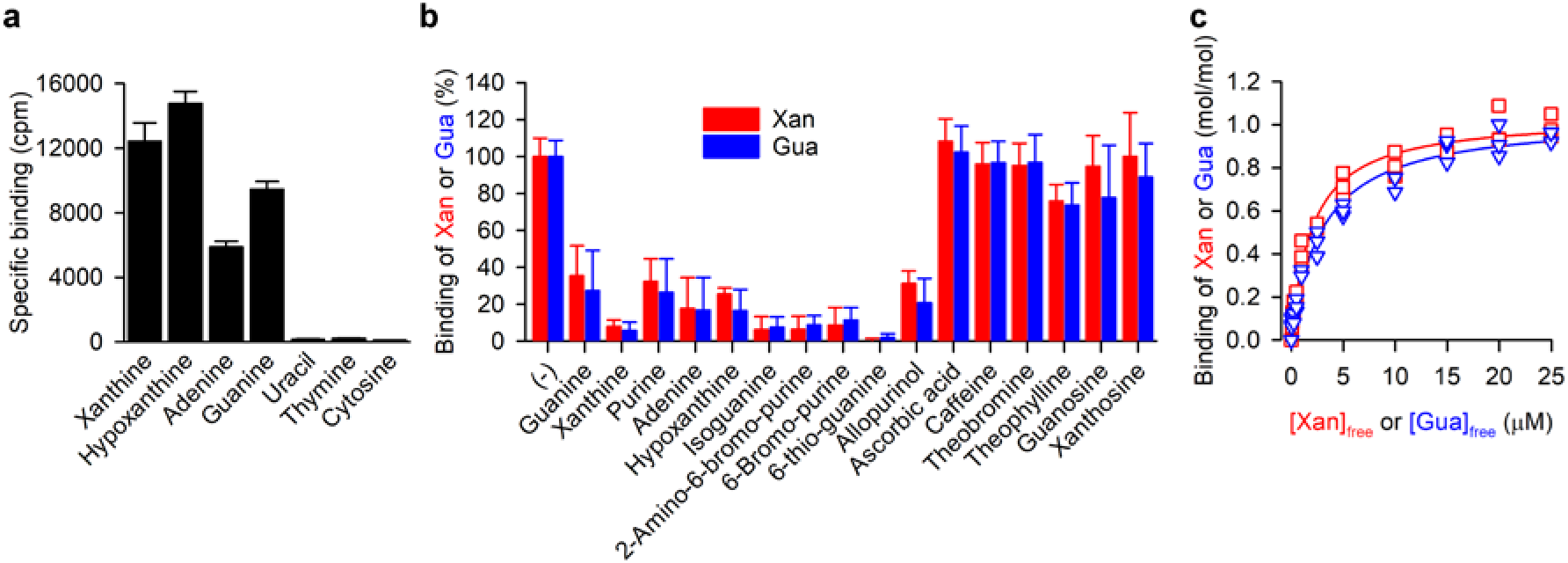
Substrate specificity of PurT_Cp_. **a.** Binding of 0.5 μM ^3^H-xanthine, ^3^H-hypoxanthine, ^3^H-adenine, ^3^H-guanine, ^3^H-uracil, ^3^H-thymine, or ^3^H-cytosine (all at a specific radioactivity of 5 Ci/mmol) was measured with the SPA using 200 ng of purified PurT_Cp_ in 200 mM Tris/Mes, pH 7.0, 10 % glycerol, 0.1 mM TCEP, 0.1 % decyl-β-D-maltopyranoside for 16 h at 4 °C. Specific binding was determined by subtracting the non-specific counts per minute (cpm) determined in the presence of 800 mM imidazole, which competes with the PurT_Cp_-His tag for binding to the Cu^2+^-His tag SPA beads, from the cpm measured in the absence of imidazole. **b.** Binding of 0.5 μM ^3^H-xanthine (red) or ^3^H-guanine (blue) to 250 ng of purified PurT_Cp_ in the presence or absence of 250 μM of the indicated compound. Data were normalized to the cpm measured in the absence of compounds for ^3^H-xanthine or ^3^H-guanine binding. Data in panels **a** and **b** are the mean±S.E.M of n≥3 (each performed as technical replicate). **c.** Saturation binding of ^3^H-xanthine (red) or ^3^H-guanine (blue) to 200 ng of purified PurT_Cp_ yielded a dissociation constant (*K_d_*) for xanthine binding of 2.02 ± 0.2 μM and a *K_d_* for guanine binding of 3.02 ± 0.33 μM with a molar substrate-to-PurT_Cp_ binding ratio of 1.04 ± 0.02 and 1.04 ± 0.03 for xanthine and guanine binding, respectively. Data of three independent experiments each performed in triplicate were subjected to global non-linear regression fitting in Prism 8.

### Structure of PurT_Cp_

The PurT_Cp_ structure was determined to a resolution of 2.80 Å and refined (R_work_/R_free_ of 21/25%). The asymmetric unit contains two protomers of PurT_Cp_ and residues 1-165 and 172-457 are resolved in each protomer (Methods and Supplementary Table S1). Both the N and C termini are located to the cytosolic side based on the positive-inside rule (22).

PurT_Cp_ forms a dimer with an extensive interface (Fig. 2a, b). Each protomer has 14 transmembrane segments (TMs), and like many other secondary solute transporters, the first half of the molecule is related to the second half by a pseudo two-fold symmetry (Fig. 2c, d). The two halves are connected by a long periplasmic loop. The 14 TMs are largely alpha helical with the exception of TMs 3 and 10, and they fold into two structurally distinct domains that we define as core and gate domains following the nomenclature used in the structure of UraA (15, 16). The gate domain is composed of TMs 5-7 from the first half and 12-14 from the second half of the protomer, and the six helices have few interactions among them and are arranged almost side by side similar to a picket fence. The gate domains from the two protomers form the dimer interface with a buried surface area of 3709 Å^2^.

**Figure 2:**
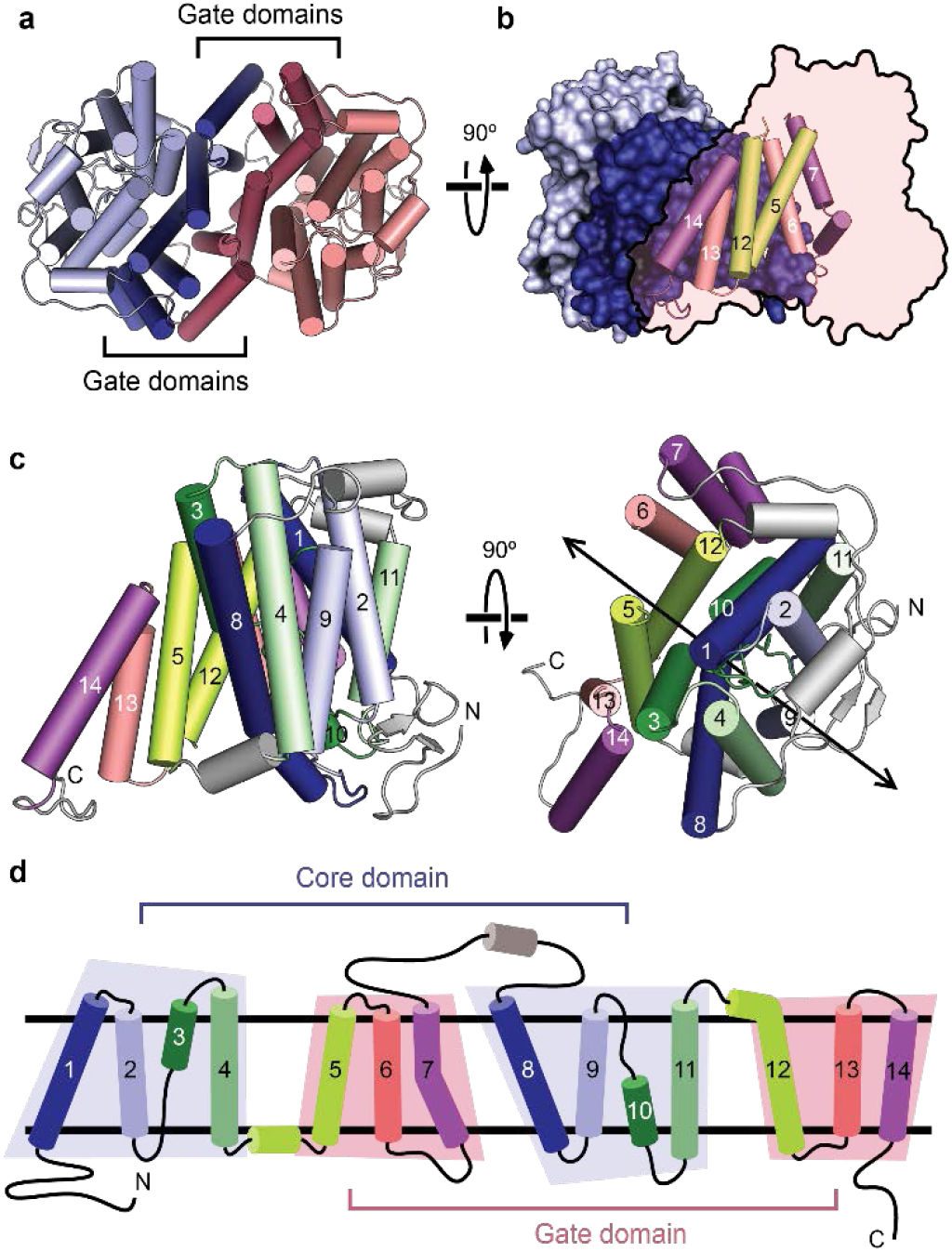
Fold and oligomeric structure of PurT_Cp_. **a.** The PurT_Cp_ dimer viewed from the extracellular side, with the gate and core domains colored in lighter or darker shades, respectively. **b.** A view of the dimer from within the plane of the membrane. One protomer is shown with a surface view, the other is shown as a transparent outline with helices from the gate domain shown as cylinders. **c.** Two perpendicular views of a protomer from the PurT_Cp_ structure colored in pairs of symmetry-equivalent transmembrane helices. The twofold pseudosymmetry axis is marked on the right. **d.** A topology diagram colored according to the same scheme as in panel **c**.

The core domain is formed by TMs 1-4 from the first half and TMs 8-11 from the second half of the protomer and the eight helices are arranged into a compact unit (Fig. 2c-d). TM 3 and its pseudosymmetry mate TM 10, are both composed of an alpha helix preceded by an extended beta strand, and the two TMs cross each other at roughly the middle of the membrane (Fig. 3a-b). The crossover region is known to bind substrates based on previous structures of the NAT family of transporters(15–17). A non-protein electron density was found near the crossover region and a guanine nucleobase was built into the density based on functional studies, although the current resolution is not sufficient to unambiguously define its identity (Supplemental Fig. 3c-d). The substrate is contained within the core domain between the ends of the half membrane-spanning helices in TM 3 and TM 10 and is occluded from the solvent on both side of the membrane (Fig. 3a-c). Two Trp residues flank the bound guanine, and potential hydrogen bonds are provided by a number of hydrophilic residues from TMs 3, 8, 10, and 12, including a highly conserved Asp residue (Asp276) on TM 8 (Fig. 3d and Supplementary Fig 4).

**Figure 3:**
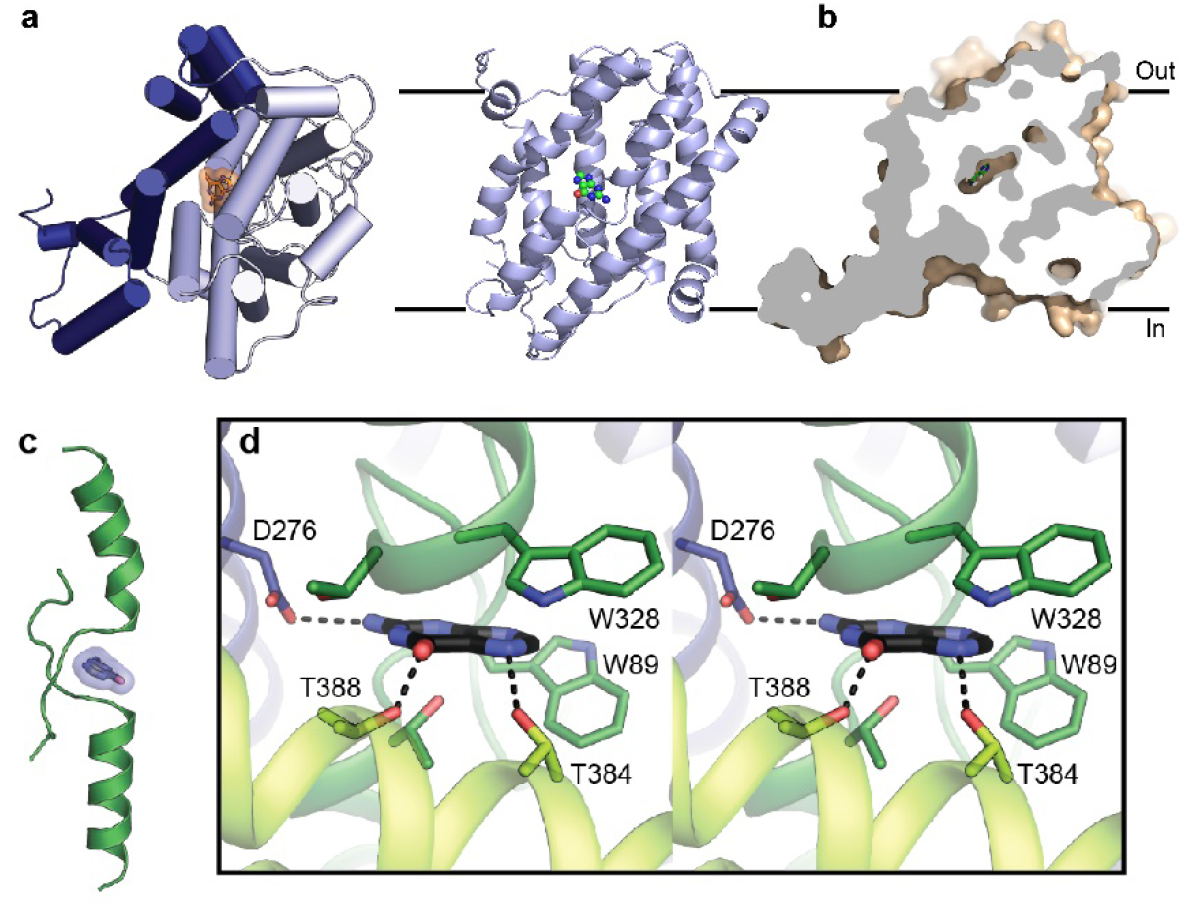
Substrate binding site. **a.** A protomer of PurT_Cp_ viewed from the extracellular side (left) and the transport domain viewed from the membrane (right), with the bound substrate molecule. **b.** Cutaway surface representation of a PurT_Cp_ monomer. **c.** The bound substrate (blue) in PurT_Cp_ is shown in relation to the TM3 and TM10 transmembrane passes. **d.** Stereo view of the substrate-binding site in PurT_Cp_, with nearby residues shown as sticks and potential hydrogen bonds marked with dashed lines.

### H^+^-coupled transport of purine in PurT_Cp_

Nucleobase transport by non-mammalian NAT members were proposed to be thermodynamically coupled to the flux of H^+^, i.e., electrogenic H^+^/nucleobase symport (23). Since residues with acidic side chains, i.e., Asp or Glu, oftentimes play central roles in the translocation of H^+^ during the symport process, we replaced conserved Asp276 with Ala or Asn to test the role of the physicochemical properties of the side chain at position 276 as potential H^+^ binding site in PurT_Cp_ and undergo protonation/deprotonation during the transport process. Binding of 0.1 μM ^3^H-xanthine or ^3^H-guanine by PurT_Cp_-D276A and -D276N was measured at pH 5.5 and pH 8.0 and compared to the binding activity by PurT_Cp_-WT. According to the ordered binding model that was proposed for the function of UraA and UapA(15, 17), H^+^ binding was proposed to occur before the binding of the co-transported substrate; consequently, increased binding of the radiolabeled purines would be expected at high H^+^ concentration (low pH). However, whereas purine binding by the two Asp276 mutants was similar to that observed for PurT_Cp_-WT at pH 8.0, lowering the pH to 5.5 almost abolished ^3^H-xanthine and ^3^H-guanine binding by the WT but did not affect the binding activity for the two Asp276 mutants (Fig. 4a). Testing the effect of the pH on ^3^H-xanthine and ^3^H-guanine binding by PurT_Cp_-WT revealed a steep pH dependence (with halfway point ~ pH 6.8) with the highest activities observed at pH ≥ 7.5 (Fig. 4b). Based on the strict pH dependence observed for binding, we tested transport of ^3^H-xanthine with PurT_Cp_-WT reconstituted into proteoliposomes of defined H^+^ gradients across the liposomal membrane. Fig. 4c shows that an inwardly directed H^+^ gradient (i.e., [H^+^]_out_ > [H^+^]_in_ or pHout < pH_in_; Δ*μ_H^+^_*) yielded the highest uptake activity. In contrast, generation of an electrical membrane potential (ΔΨ) through a valinomycin-induced K^+^ diffusion potential (inside negative) did not serve as the sole driving force for xanthine uptake, nor did it stimulate H^+^ gradient-driven xanthine accumulation. Likewise, a K^+^ diffusion potential-generated inverse membrane potential (outside negative) did not significantly affect the transport activity (Supplemental Fig. 5). Notably, xanthine uptake was impaired for PurT_Cp_-D276A and -D276N, indicating that conserved Asp276 plays a critical role in H^+^ binding and/or the regulation of substrate binding and transport by H^+^ (Supplemental Fig. 5). Testing the effect of the external pH on xanthine transport (Fig. 4d) revealed an inverse pH dependence pattern to that observed for binding (Fig. 4b). Here, the highest activity was observed at pH ≤ 5.5, and the halfway point of the curve was about pH 6.7 (Fig. 4d). Measuring the time course of ^3^H-xanthine transport (Fig. 4e) under optimized conditions (pH_in_ = 8.5, pH_out_ = pH 5.5) showed that xanthine accumulation peaked within about 5 min before falling to the concentration equilibrium (~ 1h). Determining the concentration dependence of the initial rates of uptake (measured for 10-s periods at ^3^H-xanthine concentrations between 10 nM and 25 μM) yielded a Michaelis-Menten constant (*K_m_*) of ~1.8 μM and a maximum velocity of transport (*V_max_*) of ~100 nmol x mg of PurT_Cp_^−1^ x min^−1^ (Fig. 4f). This *V_max_* translates to a catalytic turnover number (*k_cat_*) of ~0.1 s^−1^. To elucidate whether the observed pH dependence of xanthine transport is reflective of H^+^/purine symport, we tested the effect of the protonophore carbonyl cyanide 3-chlorophenylhydrazone (CCCP) on the pH-dependent transport of ^3^H-xanthine. Fig. 4g shows that dissipation of the transmembrane H^+^ gradient impairs transport by PurT_Cp_. In further support of the notion that PurT_Cp_ mediated H^+^/purine symport, solid supported membrane (SSM)-based electrophysiology using PurT_Cp_-containing proteoliposomes (pH_in_ = 8.5) showed that the addition of 10 μM xanthine to an assay medium with a pH of 5.5 elicited an inward-directed electrical current that was characterized by an initial transient component ~ 6 times larger in magnitude than that observed in control liposomes lacking PurT_Cp_ followed by a steady-state inward current that was not observed in the control liposomes. (Fig. 4h). Removal of xanthine from the assay medium elicited an inverse current transition (outside current) in PurT_Cp_-containing proteoliposomes followed by a steady-state current that was superimposable with the baseline recorded for control liposomes.

**Figure 4:**
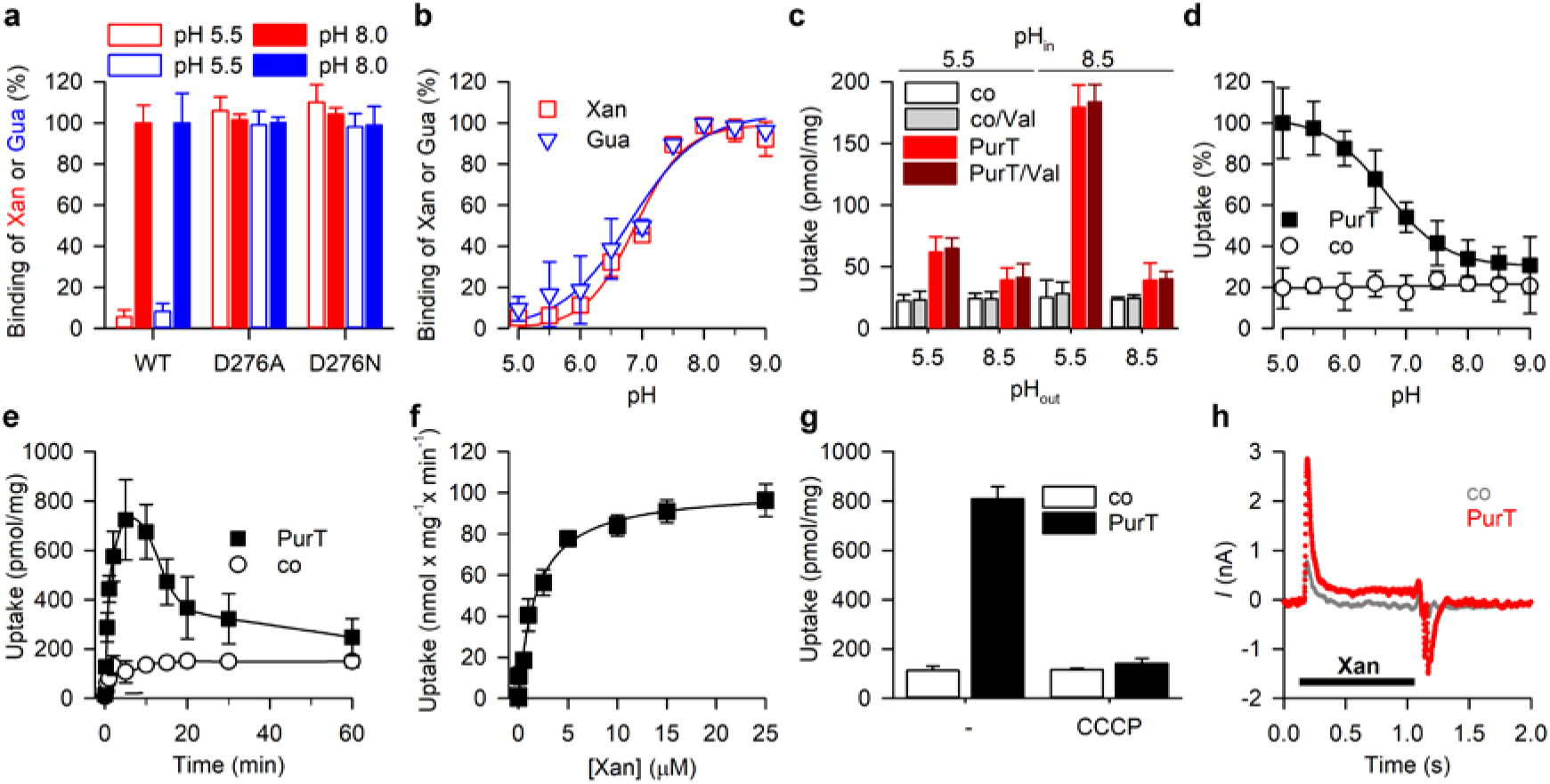
Functional characterization of PurT_Cp_. **a.** SPA-based binding of 0.5 μM ^3^H-xanthine (red) or ^3^H-guanine (blue) binding by PurT_Cp_-WT or -D276A, or D276N measured at pH 5.5 and pH 8.0. **b.** pH dependence of 0.5 μM ^3^H-xanthine or ^3^H-guanine binding by PurT_Cp_-WT. **c.** Uptake of 1 μM ^3^H-xanthine by PurT_Cp_-containing proteoliposomes requires in inwardly directed H^+^ gradient. Proteoliposomes containing PurT_Cp_ or control liposomes devoid of protein were prepared in either 20 mM Hepes-KOH, pH 8.5 or 20 mM Mes-KOH, pH 5.5 and 100 mM KCl, 2 mM β-mercaptoethanol (pH_in_) and transport was measured for 30 s in 20 mM Hepes-KOH, pH 8.5 or 20 mM Mes-KOH, pH 5.5 and 100 mM NaCl, 2 mM β-mercaptoethanol (pH_out_) in the presence or absence of 1 μM of the K^+^ ionophore valinomycin (Val) as indicated. **d.** pH dependence of 1 μM ^3^H-xanthine transport by proteoliposomes containing PurT_Cp_ or control liposomes (pH_in_ = 8.5) measured for 1 min in assay buffer with pH_out_ between 5.0 and 9.0. Data were normalized to the maximum transport activity observed for PurT_Cp_-containing proteoliposomes at pH_out_ = 5.0. **e.** Time course of 1 μM ^3^H-xanthine in proteoliposomes containing PurT_Cp_ or control liposomes (pH_in_ = 8.5; pH_out_ = 5.5). **f.** Transport kinetics of PurT_Cp_ incorporated into proteoliposomes (pH_in_ = 8.5; pH_out_ = 5.5) revealed a Michaelis-Menten constant (*K_m_*) of 1.78 ± 0.23 μM and a maximum velocity of transport (*V_max_*) of 102.0 ± 3.3 nmol x mg^−1^ x min^−1^. **g.** Substrate uptake is dependent on the proton-motive-force (pmf). Uptake of 1 μM ^3^H-xanthine in proteoliposomes containing PurT_Cp_ or control liposomes was measured for 5 min in the presence or absence of 5 μM of the protonophore carbonyl cyanide 3-chlorophenylhydrazone (CCCP). Data in panels **a** – **g** are means ± S.E.M. of three independent experiments performed as technical triplicates. **h.** Transport of xanthine is accompanied by H^+^ flux. Representative recording (n=8) of a solid supported membrane (SSM) electrophysiological measurement performed with proteoliposomes containing PurT_Cp_ (red) or with control liposomes (grey) using the SURFE^2^R N1 (Nanion Technologies). 10 μM xanthine were added to the assay buffer as indicated by the bar.

## Discussion

The structure of PurT_Cp_, together with functional data presented here provides further insight into the underlying elements of ion-coupled substrate transport by the H^+^-dependent members of the NAT family. The purine binding site is located to the crossover region of TM3 and 10, and this feature is conserved in transporters of the same structural fold (Supplemental Fig. 6). The PurT_Cp_ structure was captured in an occluded conformation in which the substrate does not have unimpeded access to either side of the membrane.

Our functional data support H^+^/substrate symport as the underlying mechanism of nucleobase transport by PurT_Cp_. While the transport of nucleobases is strictly dependent on the presence of a transmembrane H^+^ gradient (Δ*μ_H^+^_*), SSM-based electrophysiological measurements employing PurT_Cp_-containing proteoliposomes reveal a change in the membrane potential as a result of H^+^ influx into the proteoliposomes concomitant with the translocation of nucleobases. This notion is supported by the finding that the protonophore CCCP dissipates PurT_Cp_-mediated nucleobase transport. Our functional data further highlight the essential character of conserved Asp276, a residue that was identified to interact with the bound guanine (the carboxy oxygen of Asp276 binds the guanine amino group) in the PurT_Cp_ structure. H^+^-coupled nucleobase transport was impaired with PurT_Cp_ variants in which Asp276 was replaced with Ala or Asn. However, in contrast to PurT_Cp_-WT, binding of nucleobases by these variants was pH independent, thus hinting to a transport mechanism that involves Asp276 in the coordinated association and dissociation of the symported H^+^. Under the equilibrium binding conditions in the SPA (i.e., the lack of Δ*μ_H^+^_* across the membrane; i.e., the extra- and intracellular face of PurT_Cp_ are exposed to the same pH), nucleobase binding by purified PurT_Cp_-WT decreased with increasing H^+^ concentrations, suggesting that protonation of Asp276 interferes with nucleobase binding under those conditions. This notion is supported by the fact that PurT_Cp_ variants with amino acid side chains at position 276 that cannot be protonated feature virtually indistinguishable nucleobase binding activities at all pH values tested. Since these Asp276 mutants fail to mediate H^+^-coupled nucleobase symport, it appears possible that protonation of Asp276 in PurT_Cp_-WT is the trigger for conformational changes that coordinate the dissociation of bound substrate and H^+^ from their respective binding sites to the cytoplasmic side of PurT_Cp_. It is thus feasible to speculate that H^+^ binding to Asp276 may induce conformational changes to nucleobase-bound PurT_Cp_ that lead to the release of H^+^ and nucleobase substrate in the cytoplasm. This model supports the common paradigm of the alternating access mechanism according to which the binding of the coupling cation induces a conformation that is required for the coordinated translocation of the substrate molecule across the membrane.

## Methods

### Expression, purification and crystallization

100 bacterial homologs of nucleobase/cation symporter 2 (NCS2) proteins were cloned into a modified pET28 plasmid (Novagen) with either an N- or a C-terminal deca-histidine tag and a TEV protease recognition site at the central facility of the New York Consortium on Membrane Protein Structure (NYCOMPS) as described in Love *et al*. (24). Expression vectors containing gene of interest were transformed into BL21-Gold (DE3) competent cells (Agilent) and the cells were cultured to a density of ~1 OD_600_/ml at 37 °C. Overexpression of protein was then induced by addition of IPTG to a final concentration of 0.5 mM at 20 °C for overnight. For expression screening, 10 ml of cell cultures were harvested, and cell pellets were resuspended in 1 ml of lysis buffer containing 20 mM HEPES pH 7.5, 150 mM NaCl, 10% (v/v) glycerol, 2 mM β-mercaptoethanol and 1 mM PMSF. Cells were lysed by sonication and the protein was extracted with 30 mM n-dodecyl-β-D-maltopyranoside (DDM) in the lysis buffer for 2 h at 20 °C. The cell lysate was centrifuged at 40,000 × *g* for 45 min at 4 °C and the supernatant was loaded onto a cobalt affinity column (Clontech). After washing the column with 20 bed volume of 20 mM HEPES pH 7.5, 150 mM NaCl, 10% (v/v) glycerol, 2 mM β-mercaptoethanol, 5 mM DDM, 20 mM imidazole pH 8.0, bound protein was eluted in the same washing buffer except with 300 mM imidazole pH 8.0. Protein samples were then subjected to SDS-PAGE and Commassie staining. Expressed clones were scaled-up to 2-12 liter cultures and purified proteins were analyzed by size-exclusion chromatography. One clone from *Colwellia psychrerythraea* 34H (PurT_Cp_) with an N-terminal deca-histidine tag yielded ~1 mg protein/liter culture and displayed a mono-dispersed peak when eluted from a size-exclusion column, representing a suitable candidate for crystallization trials.

Purification of PurT_Cp_ was modified slightly from the above described as the last elution step was omitted. Instead, PurT_Cp_ bound on a cobalt column was released by TEV protease cleavage for 1 hr at 20 °C to separate cleaved PurT_Cp_ from uncleaved protein. Purified PurT_Cp_ was then loaded onto a Superdex 200 10/300 GL column (GE Health Sciences) equilibrated in 20 mM HEPES pH 7.5, 150 mM NaCl, 5 mM β-mercaptoethanol, 12 mM n-nonyl-β-D-maltopyranoside (NM) and 4 mM 3-([3-cholamidopropyl]dimethylammonio)-2-hydroxy-1-propanesulfonate (CHAPSO).

### Crosslinking

Purified PurT_Cp_ or UraA (~ 1 mg/ml) was incubated with indicated concentrations of disuccinimidyl glutarate (DSG) at room temperature for 10 min and loaded onto SDS-PAGE. Protein bands were visualized by Commassie staining.

### Crystallization

Purified PurT_Cp_ protein was concentrated to ~10 mg/ml as approximated by ultraviolet absorbance and crystallization was set up using the sitting-drop vapor diffusion method. The best PurT_Cp_ crystals were grown at 4 °C in 30% (v/v) PEG-400, 0.1 M MES pH 6.0, 3 mM Na-glycochenodeoxycholate and 0.5 mM 6-bromopurine and reached full size within 7 days. The crystals were cryoprotected by raising PEG-400 (v/v) concentrations gradually to 38 % with an increment of 2 % over a course of 16 h before being flash frozen in liquid propane.

### X-ray data collection and processing

X-ray data were collected at xxx at the Advanced Photon Source at Argonne National Laboratory. Native dataset was collected at a wavelength of 0.9150 Å with a resolution of 2.85 Å. Anomalous dataset was collected at a wavelength of xxx Å with a resolution of xx Å. The reflections were then processed with HKL2000 (25). The crystal belonged to spacegroup P21212 (No.18) with unit cell dimensions of 131.713 Å, 135.993 Å, and 79.169 Å.

### Structure determination and refinement

The initial phase was determined by molecular replacement using the UraA structure (PDB ID 5XLS) as the searching model. The structure was completed through successive rounds of model building in COOT (26) and refinement in PHENIX (27). Protein geometry was validated with MolProbity(28).

### Scintillation Proximity Assay

Binding of nucleobases was performed by means of the scintillation proximity assay (SPA) using His-tagged PurT_Cp_ in conjunction with Copper HIS-Tag YSI beads (Perkin Elmer, RPNQ0096). 250 ng of purified PurT_Cp_ were immobilized on 125 μg SPA beads per 100-μL assay in 200 mM Tris/Mes at the indicated pH, 20 % (v/v) glycerol, 1 mM TCEP, 0.1 % DDM. ^3^H-labeled xanthine (5.2 Ci/mmol), hypoxanthine (9 Ci/mmol), adenine (29.7 Ci/mmol), guanine (21.2 Ci/mmol), or uracil (38.7 Ci/mmol) (all radiochemicals were purchased from American Radiolabeled Chemicals, Inc.).

### Transport measurements

Purified PurT_Cp_-WT, -D276A, or -D279N was reconstituted into pre-formed liposomes made of total *E. coli* lipids (Avanti) at a 1:100 (w/w) ratio as described(29). Liposomes were prepared in *i*) 20 mM Hepes-KOH, pH 8.5, 100 mM KCl, 2 mM β-mercaptoethanol, *ii*) 20 mM Mes-KOH, pH 5.5, 100 mM KCl, 2 mM β-mercaptoethanol, *iii*) 20 mM Hepes-KOH, pH 8.5, 100 mM NaCl, 2 mM β-mercaptoethanol, or *iv*) 20 mM Mes-KOH, pH 5.5, 100 mM NaCl, 2 mM β-mercaptoethanol. The same buffers (in all possible cis/trans permutations) were also used as assay buffer to test the influence of the membrane potential on the transport reaction, by generating a valinomycin-mediated K^+^ diffusion potential. To test the effect of the external pH on the uptake activity, the external buffer was composed of 20 mM Mes-KOH, pH 5.5 - 6.5 or 20 mM Hepes-KOH, pH 7.0 – 9.0 and 100 mM KCl, 2 mM β-mercaptoethanol. Uptake of ^3^H-xanthine was measured with a rapid filtration assay using 0.45 μM nitrocellulose filters (Millipore). Reactions were incubated at 23 °C for the indicated periods of time and quenched by the addition of ice-cold 100 mM potassium phosphate, pH. 6.0, 100 mM LiCl before filtration. The radioactivity retained on the filters was determined with scintillation counting using the dried filters. Known amounts of radioactivity were used to convert decays per minute (dpm) to mol.

### Microscale Thermophoresis

Binding of either guanine or xanthine by purified PurT_Cp_ was measured by microscale thermophoresis (MST) using the Monolith NT.LabelFree and Monolith NT.115 (NanoTemper Technologies, Germany) to detect the thermophoretic signals of native tryptophan and tagged fluorescence in PurT_Cp_, respectively. For label-free binding measurements, 500 nM PurT_Cp_ were mixed with guanine or xanthine at a concentration range of 0.47 – 94 μM in MST assay buffer containing 20 mM HEPES, pH 7.5, 150 mM NaCl, 5 mM MgCl_2_, 10 % v/v glycerol, 2 mM β-mercaptoethanol and 0.1% w/v DM (n-decyl-β-D-maltopyranoside). The mixture was incubated at room temperature for 20 min and then loaded into Monolith NT.LabelFree capillaries. Measurements were conducted at high (60 %) MST power and 15 % excitation power using the MO.Control v1.4.4 software. For binding assays using the Monolith NT.115, prior to the MST measurement, purified PurT_Cp_ was labeled with RED fluorescent dye NT-647 (RED-tris-NTA; NanoTemper Technologies) following the manufacturer’s protocol. As for label-free measurements, assays were performed using 50 nM RED-labeled PurT_Cp_ in MST assay buffer in Monolith NT.115 Premium capillaries and the thermophoresis reaction was set at high MST power. MST raw data, after evaluation in MO.Control, were transferred, normalized, and analyzed using non-linear regression fitting in GraphPad Prism 8.

#### Solid supported membrane (SSM) electrophysiology

SSM electrophysiological measurements were performed using the SURFE^2^R N1 (Nanion Technologies) according to published protocols(30). Briefly, the sensors were filled with 1.5 μL of the lipid solution (1,2-diphytanoyl-sn-glycero-3-phosphocholine in n-decane), 50 μL of non-activating buffer (20 mM MES, pH 5.5, 100 mM NaCl) and 10 μL of PurT_Cp_-containing proteoliposomes (after being extruded and sonicated). The activating solution (20 mM MES, pH 5.5, 100 mM NaCl, 10 μM xanthine) was applied with the single-solution exchange protocol (activating buffer incubation for 1 s). Four different data sets from individual sensors were recorded. Peak currents were corrected by subtracting the peak currents recorded with control liposomes (devoid of PurT_Cp_). To obtain the PurT_Cp_-elicited charge (Coulomb) movement associated with xanthine transport, the area under the curve (current as function of time) was analyzed with GraphPad Prism 8.

## Accession Codes

Atomic coordinates and structure factors have been deposited with the Protein Data Bank under accession ID 7TAK.

## Acknowledgements

This work was supported by the US National Institutes of Health grant R01 GM119396, DK122784, and HL086392.

