## Supplementary Information for "Insight into the mechanism of H^+^-coupled nucleobase transport"

### Supplementary Figures

#### Supplementary Figure 1

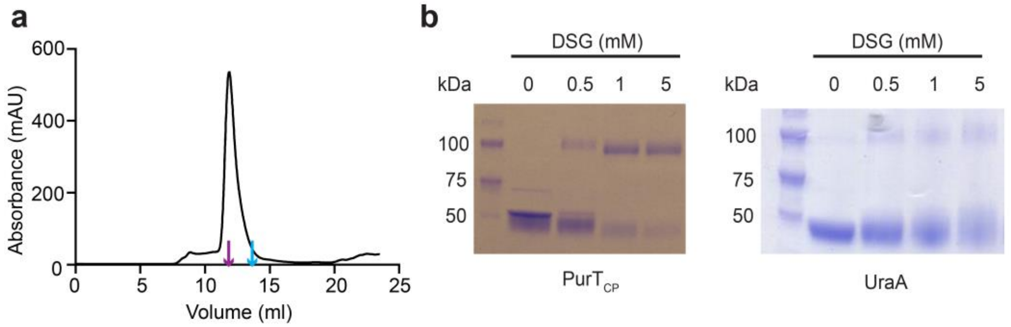


**Supplementary Figure 1. Purification and crosslinking of PurT$_{Cp}$. a.** Size-exclusion chromatography of PurT$_{Cp}$ in DDM. Elution volumes of membrane proteins of known molecular weight, bcMalT (100 kDa, purple) (1) and mouse SCD1 (41 kDa, blue) (2) are marked by arrows. **b-c**. SDS-PAGE showing Crosslinking of purified PurT$_{Cp}$ (**b**) or UraA (**c**) by DSG.

**Supplementary Figure 2**

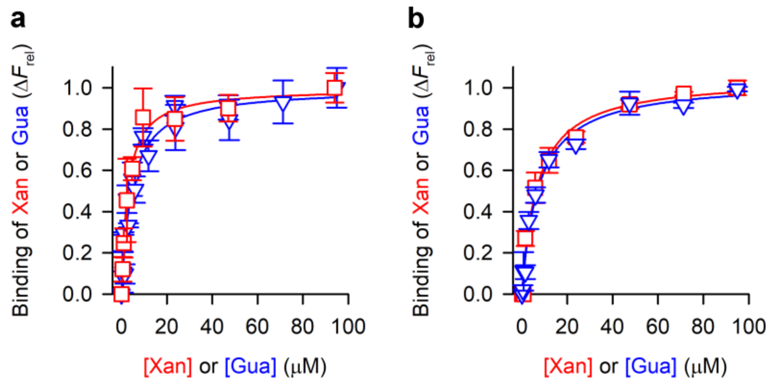


**Supplementary Figure 2: Substrate binding kinetics of PurT$_{Cp}$.** Saturation binding of xanthine (red) or guanine (blue) to 0.5 µM of purified PurT$_{Cp}$ was measured with microscale thermophoresis (MST) using **a.** the Monolith NT.LabelFree in conjunction with unlabeled PurT$_{Cp}$ or **b**. the Monolith NT.115 in combination with RED-tris-NTA-labeled PurT$_{Cp}$. Data of ≥ 6 independent experiments were subjected to non-linear regression fitting in Prism 8 and yielded a $K_d$ for xanthine binding of 2.76 ± 0.4 µM and a $K_d$ for guanine binding of 4.47 ± 0.75 µM with the label-free method (panel **a**). Fitting of data measured with the RED NT-647 dye-labeled PurT$_{Cp}$ yielded a $K_d$ for xanthine binding of 6.81 ± 1.25 µM and a $K_d$ for guanine binding of 7.37 ± 0.93 µM.

**Supplementary Figure 3**

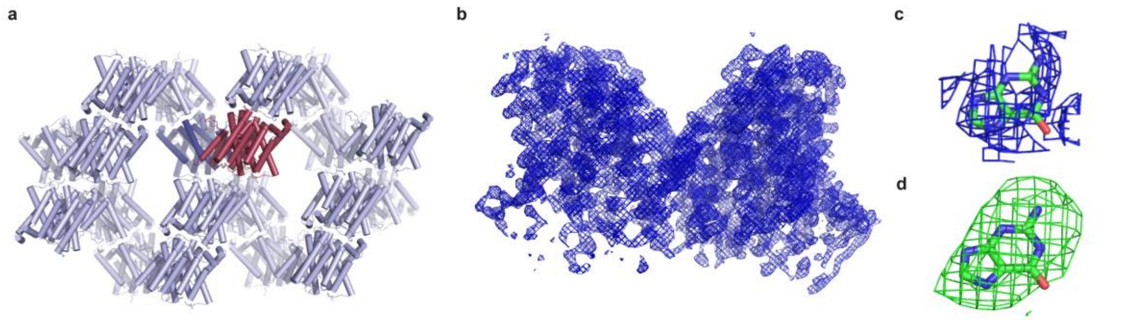


**Supplementary Figure 3. Crystal packing and electron density of PurT$_{Cp}$. a.** A cross-section of the crystal lattice in the UraA structure. One asymmetric unit is colored pink. **b.** $2F_O$-$F_C$ map of a PurT$_{Cp}$ dimer contoured at 2 σ**. c**. $2F_O$-$F_C$ map of a guanine molecule contoured at 1.0 σ. **d**. $F_O$-$F_C$ map of a guanine molecule contoured at 2.5 σ.

**Supplementary Figure 4**

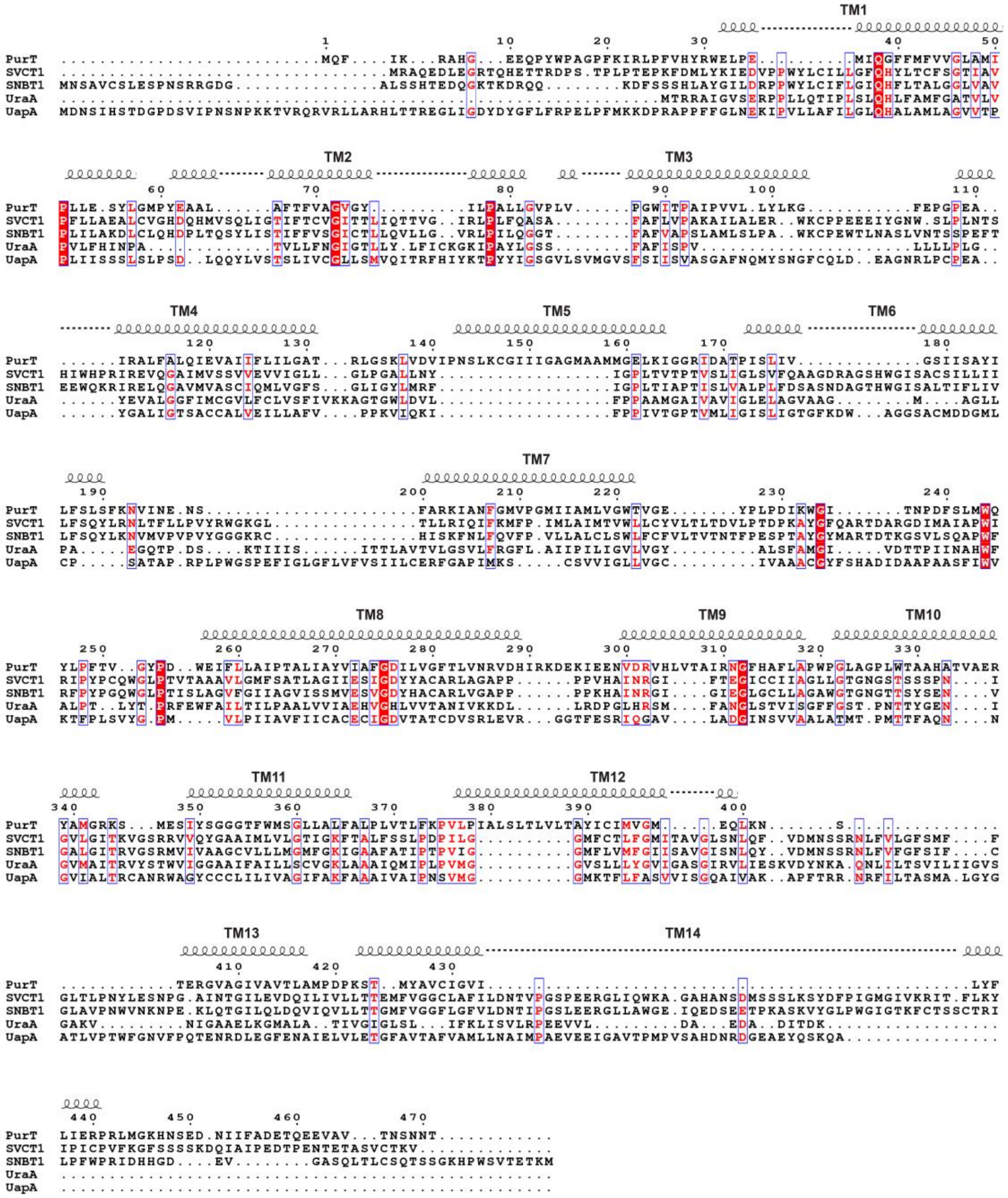


**Supplementary Figure 4. Sequence alignment of PurT with selected NAT family members.** PurT (Gene Bank accession number OUR80909) from *Colwellia psychrerythraea* 34H; SVCT1 (Uniprot accession number BC050261), a human L-ascorbate transporter; SNBT1 (AB511909), an uracil transporter from rat; UraA (YP490725), an uracil transporter from *E. coli;* UapA (Q07307), a uric acid/xanthine $H^+$ symporter from *A. nidulans*. The selected sequences were aligned using Clustal Omega1 and plotted using Espript server. The secondary structure elements of PurT are indicated above the sequence. $PurT_{Cp}$ shares 12 % identity and 25.3 % similarity with human SVCT1 (SLC23A1), 12.9 % identity and 26.4 % similarity with rat SVCT1, 16.4 % identity and 31.4 % similarity with *E. coli* UraA, and 13.2 % identity and 25.6 % similarity with *A. nidulans* UapA.

**Supplementary Figure 5**

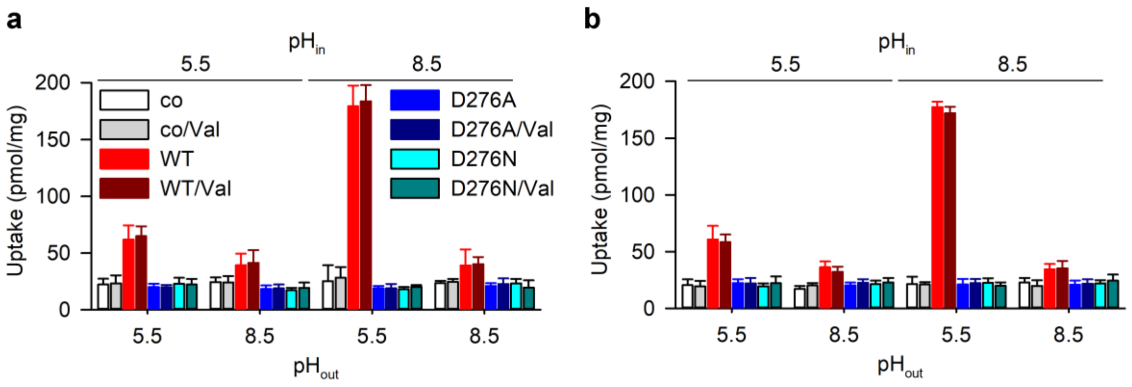


**Supplementary Figure 5.** Characterization of PurT$_{Cp}$-mediated transport. **a.** Effect of an inwardly-directed membrane potential. Uptake of 1 µM $^3$H-xanthine by proteoliposomes containing PurT$_{Cp}$-WT, -D276A, or -D279N, or control liposomes prepared in 20 mM Hepes-KOH, pH 8.5, 100 mM KCl, 2 mM β-mercaptoethanol (pH$_{in}$ = 8.5) or 20 mM Mes-KOH, pH 5.5, 100 mM KCl, 2 mM β-mercaptoethanol (pH$_{in}$ = 5.5) was measured for 30 s in 20 mM Hepes-KOH, pH 8.5, 100 mM NaCl, 2 mM β-mercaptoethanol (pH$_{out}$ = 8.5) or 20 mM Mes-KOH, pH 5.5, 100 mM NaCl, 2 mM β-mercaptoethanol (pH$_{out}$ = 5.5) in the presence or absence of 1 µM of valinomycin (Val) as indicated. Under this condition, the addition of the potassium-selective ionophore valinomycin generated an outwardly-directed flux of $K^+$ that leads to the generation of an inwardly-directed electrical membrane potential (inside negative). **b.** Effect of an outwardly-directed membrane potential. Uptake of 1 µM $^3$H-xanthine by proteoliposomes containing PurT$_{Cp}$-WT, -D276A, or -D279N or control liposomes (bar fills are identical to those shown in panel **a.**) prepared in 20 mM Hepes-KOH, pH 8.5, 100 mM NaCl, 2 mM β-mercaptoethanol (pH$_{in}$ = 8.5) or 20 mM Mes-KOH, 100 mM NaCl, 2 mM β-mercaptoethanol (pH$_{in}$ = 5.5) was measured for 30 s in 20 mM Hepes-KOH, pH 8.5 (pH$_{out}$ = 8.5), 100 mM KCl, 2 mM β-mercaptoethanol or 20 mM Mes-KOH, pH 5.5, 100 mM KCl, 2 mM β-mercaptoethanol (pH$_{out}$ = 5.5) in the presence or absence of 1 µM of valinomycin (Val) as indicated. Here, the addition of valinomycin mediated the influx of $K^+$, leading to the generation of an outwardly-directed membrane potential (inside positive). Data are means ± S.E.M. of ≥ 3 independent experiments performed as technical triplicates.

**Supplementary Figure 6**

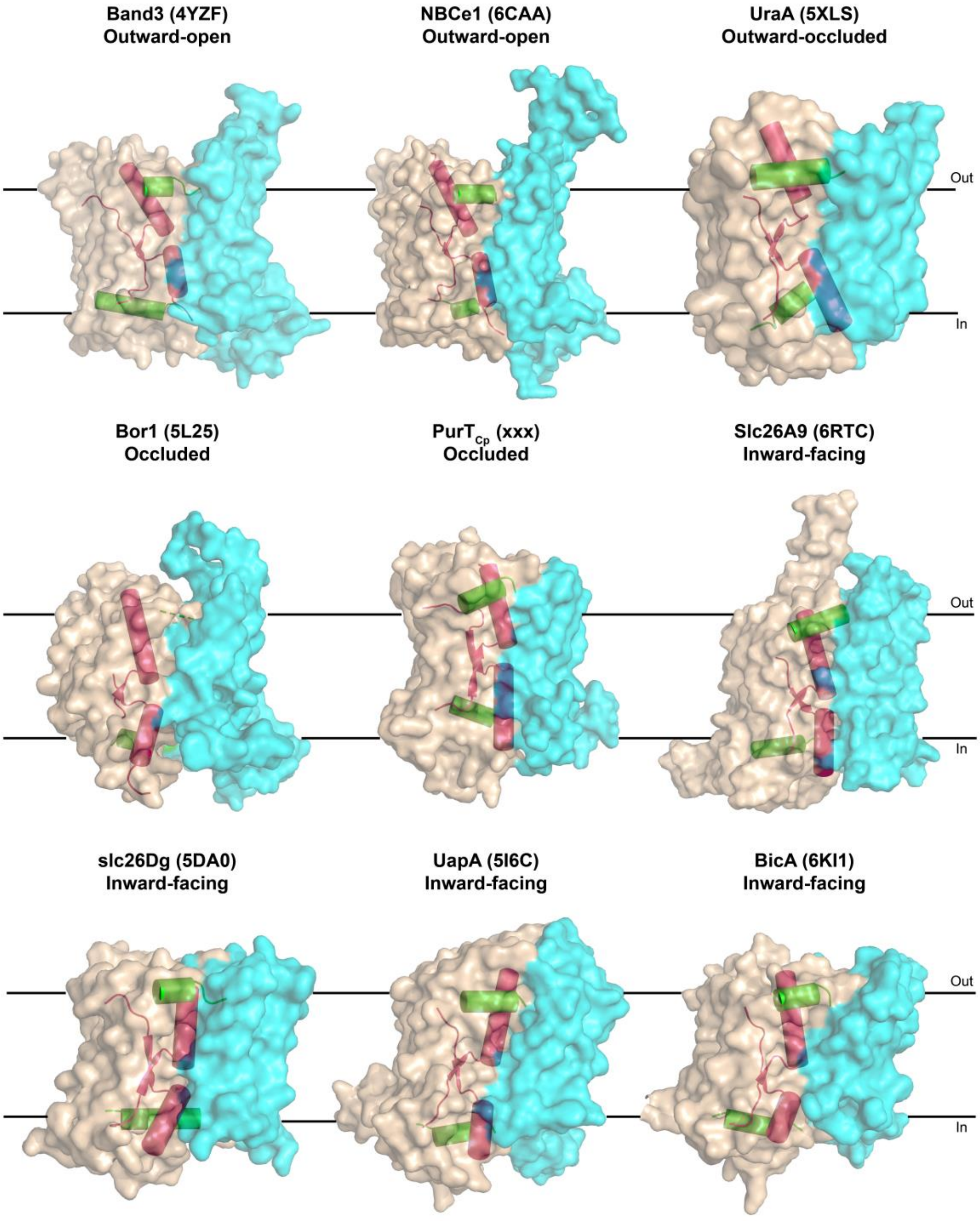


**Supplementary Figure 6. Conformations of proteins with similar structural fold to PurT$_{Cp}$.** One protomer of each protein is shown as cartoon representation. The interface domains are colored cyan and the transport domains wheat. TM3 and TM10 that form the substrate binding site are highlighted as red cylinders. The two amphipathic helices that bridge the interface domain and the transport domain are shown as green cylinders.

**Supplementary Table**

### Table S1. Data collection and refinement statistics

| *Dataset* | Native $PurT_{Cp}$ |
|---|---|
| **Data Collection** | |
| Space group | P21212 (No. 18) |
| Unit cell (Å) | a=131.71, b=135.99 c=79.17 |
| Wavelength (Å) | 0.9150 |
| Resolution (Å) | 50.00 – 2.75 (2.80 – 2.75) |
| Completeness (%) | 98.63 (90.43) |
| Redundancy | 7.5 (6.1) |
| Mean I/σ | 43.18 (2.17) |
| **Refinement** | |
| Resolution (Å) | 41.78 – 2.80 (2.90 – 2.80) |
| Unique reflections | 35339 (3186) |
| $R_{work}$ (%) / $R_{free}$ (%) | 21.30/24.59 |
| Number of non-hydrogen atoms | |
| Protein | 6847 |
| Ligand/ion | 22 |
| Average B factors ($Å^2$) | |
| Protein | 105.70 |
| Ligand/ion | 104.86 |
| Rmsd | |
| Bond length (Å) | 0.008 |
| Bond angles (°) | 1.19 |
| Ramachandran plot (%) | |
| Favored | 97.06 |
| Allowed | 2.94 |
| Outliers | 0.00 |
| Rotamer outliers (%) | 0.69 |

Statistics for the highest-resolution shell are shown in parentheses.
